# CRISPR perturbations at many coronary artery disease loci impair vascular endothelial cell functions

**DOI:** 10.1101/2021.02.10.430527

**Authors:** Florian Wünnemann, Thierry Fotsing Tadjo, Melissa Beaudoin, Simon Lalonde, Ken Sin Lo, Guillaume Lettre

**Affiliations:** Montreal Heart Institute, Montréal, Québec, Canada; Faculté de Médecine, Université de Montréal, Montréal, Québec, Canada

**Author notes:** **Correspondence:** Guillaume Lettre, Montreal Heart Institute, 5000 Belanger Street, Montreal, Quebec, Canada, H1T 1C8.

## Abstract

Genome-wide association studies have identified 161 genetic variants associated with coronary artery disease (CAD), but the causal genes and biological pathways remain unknown at most loci. Here, we used CRISPR knockout, inhibition and activation to target 1998 variants at 83 CAD loci to assess their effect on six vascular endothelial cell phenotypes (E-selectin, ICAM1, VCAM1, nitric oxide, reactive oxygen species, calcium signalling). We identified 42 significant variants located within 26 CAD loci. Detailed characterization of the RNA helicase *DHX38* and CRISPR activation at the *FURIN/FES, CCDC92/ZNF664* and *CNNM2* loci revealed a strong effect on vascular endothelial cell senescence.

## INTRODUCTION

Coronary artery disease (CAD) remains the main cause of mortality in the world despite widely available drugs (*e.g.* statins) and the known benefits of simple prevention strategies (*e.g.* exercise). Part of the complexity to prevent and treat CAD resides in our incomplete understanding of atherosclerosis, the pathophysiological process largely responsible for CAD initiation and progression. Atherosclerosis is triggered by many environmental risk factors and other intrinsic stimuli, and results in the dysregulation of vascular wall homeostasis due to the accumulation of cholesterol-rich lipoproteins and a maladaptive inflammatory state^1,2^.

Human genetics provide a framework to dissect the biological pathways and cellular networks implicated in atherosclerosis. Genome-wide association studies (GWAS) have already identified 161 loci associated with CAD^3,4^. However, the functional characterization of genes that modulate CAD risk at GWAS loci is labor-intensive. It is further complicated by the fact that most CAD variants are non-coding and are in linkage disequilibrium (LD) with a multitude of other DNA sequence variants.

Half of the CAD GWAS loci do not associate with traditional risk factors. We and others have hypothesized that some of the CAD variants, which are enriched in open chromatin regions found in human vascular endothelial cells, directly modulate endothelial cell functions^5–9^. The functional characterization of two CAD genes in endothelial cells, *PLPP3*^10^ and *MIA3/AIDA*^9^, has further supported this hypothesis. Vascular endothelial cells have critical roles in atherosclerosis^9,11,12^. Upon activation, they express adhesion molecules necessary for monocyte rolling and attachment (*e.g.* E-selectin, ICAM1, VCAM1) and weakening of their cell-cell junctions can facilitate monocyte transmigration into the intima. Furthermore, dysfunctional endothelial cells adopt an atheroprone behaviour with changes in calcium (Ca^2+^) signalling^13^, decreased bioavailability of the vasodilator nitric oxide (NO) and increased production of reactive oxygen species (ROS).

The development of pooled CRISPR-based screens now allows perturbation experiments to test most sentinel and LD proxy variants associated with CAD for a role in human vascular endothelial cells^14^. Moreover, by using inhibition (KRAB) or activation (VP64) domains tethered to an inactivated Cas9 (dCas9), it is possible to mimic loss- or gain-of-function effects that might elude perturbations due to classic Cas9 insertion-deletions (indels)^15–17^. Here, we carried out comprehensive pooled CRISPR screens for six endothelial phenotypes relevant to atherosclerosis (presentation of adhesion proteins at the cell membrane (E-selectin, ICAM1 and VCAM1), production of NO and ROS, and intracellular Ca^2+^ concentration) using three different Cas9 perturbation modalities (double-strand break induction (Cas9), inhibition (dCas9-KRAB or CRISPRi) and activation (dCas9-VP64 or CRISPRa)). Our results identified 42 variants that modulate endothelial functions, including a subset that cause endothelial dysfunction by inducing vascular endothelial cell senescence.

## RESULTS

### FACS-based pooled CRISPR screens for endothelial functions

The design of our sgRNA library is summarized in **Figure 1A**. To target genomic regions associated with CAD, we collected 92 GWAS sentinel variants at 89 CAD-associated loci^5,6,18,19^ and retrieved their proxy variants in strong LD (*r*^2^ >0.8 in populations of European ancestry). Using this strategy, we derived a set of 2,893 variants (92 GWAS sentinel and 2,801 LD proxy variants) (**Figure 1A** and **Supplementary Table 1**). For each of these variants, we designed a maximum of five high-quality sgRNAs (**Figure 1B**). The mean distance between sgRNA potential cut-sites and the targeted variants was 22-bp (**Figure 1C**). After quality-control steps, we generated a list of 7,393 sgRNA that targeted 1,998 variants at 83 CAD loci (**Supplementary Table 2**). On average at each CAD locus, our sgRNA library covered 76±22% of the targeted variants (**Figure 1D** and **Supplementary Table 3**). Of the 83 tested loci, we could capture 100% of the targeted variants at 20 CAD loci and ≥80% of variants at 38 loci (**Supplementary Table 3**). The majority of the targeted variants were intronic (70.8%) or intergenic (10.2%) (**Figure 1E**)^20,21^.

**Figure 1.**
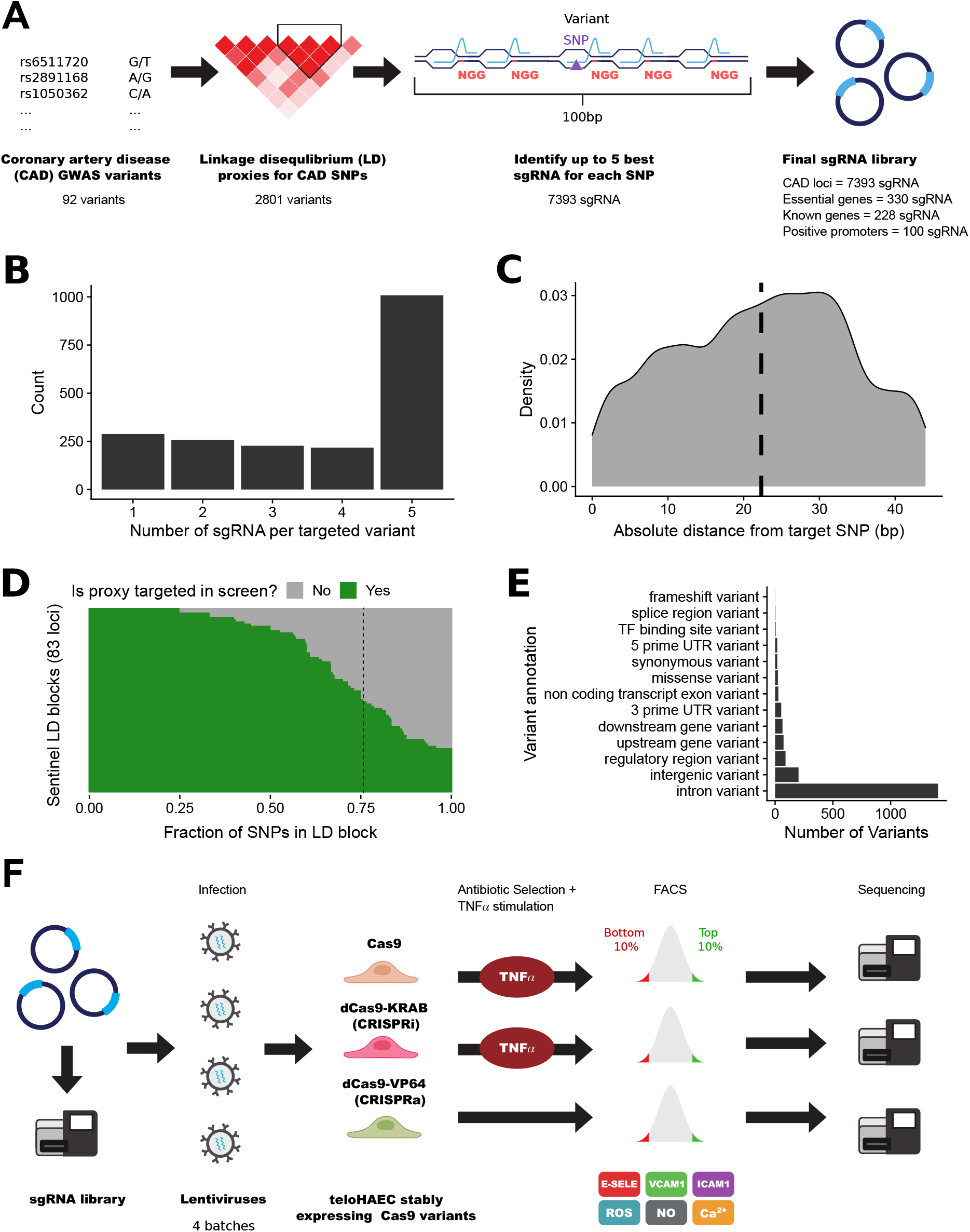
Pooled CRISPR screens to identify CAD variants and genes that modulate vascular endothelial functions. (**A**) From 92 loci associated with coronary artery disease (CAD) risk by genome-wide association studies (GWAS), we identified 2893 sentinel and linkage disequilibrium proxy variants for testing. For each of these variants, we attempted to design a maximum of five high-quality guide RNAs (sgRNAs) within a 100-bp window. In the design of the library, we also included sgRNAs that target genes essential for cell viability, as well as sgRNAs that target the coding sequence and promoter of genes that control endothelial cell functions (known genes, positive controls). (**B**) Number of sgRNAs per targeted variant that passed stringent quality-control filters. In total, we designed 7393 sgRNAs against 1998 CAD-associated variants (mean and median number of sgRNA per variant are 3.7 and 5, respectively). (**C**) Distribution of the absolute distance of the sgRNA cut-site relative to the targeted variant in base pairs (the vertical dashed line indicates mean sgRNA distance). (**D**) Fraction of variants at each locus that are successfully targeted by our pooled CRISPR screens. Each row represents one of the CAD loci that we tested. In green is the fraction of variants - including sentinel and LD proxies - for which we designed high-quality sgRNAs and obtained results for the endothelial function phenotypes. On average, 76% of variants at any given CAD locus are captured in the screens (vertical dashed line). (**E**) Most severe annotation for the 1998 CAD variants targeted by the lentiviral sgRNA libraries using ENSEMBL’s Variant Effect Predictor (VEP) module. (**F**) As a control step, we sequenced the plasmid library to ensure even representation of sgRNAs in the pool. Then, we produced four independent batches of lentiviruses which we used to infect teloHAEC cells that stably express Cas9, dCas9-KRAB (CRISPRi) or dCas9-VP64 (CRISPRa). Following antibiotic selection and TNFα treatment (for Cas9 and CRISPRi), we stained teloHAEC for cell surface markers (E-selectin, ICAM-1, VCAM-1) or intracellular signaling molecules (reactive oxygen species (ROS), nitric oxide (NO), calcium (Ca^2+^)). By flow cytometry, we sorted cells from the bottom and top 10 percentiles of the marker distributions, and sequenced sgRNAs found in each fraction.

We utilized lentiviruses to deliver our pooled CRISPR libraries to immortalized human aortic endothelial cells (teloHAEC) that stably express one of three Cas9 variants (Cas9, CRISPRi, CRISPRa) (**Figure 1F**). We treated Cas9 and CRISPRi (but not CRISPRa) infected cells with TNFα in order to find genes that can block (Cas9, CRISPRi) or induce (CRISPRa) a pro-inflammatory response. Then, we labelled cells with fluorescent antibodies against E-selectin, VCAM1, or ICAM1, or with fluorescent dyes for signalling molecules (ROS, NO, Ca^2+^), and sorted cell populations by flow cytometry (FACS) to collect the bottom and top 10% cells based on fluorescence intensity (**Figure 1F** and **Supplementary Figures 1-3**). We amplified and sequenced the sgRNAs from the FACS cell fractions to identify sgRNAs that have a significant effect on endothelial functions. Quality-control analyses of sorted cell fractions showed a good representation of sgRNA diversity (mean Gini index=0.076±0.01) and a good read coverage per sgRNA (mean number of aligned reads per sgRNA=1995±2981) (**Supplementary Figure 4**). Analysis of the 10% most variable sgRNAs across all assays revealed clustering of samples along the Cas9 modalities (**Figure 2A**).

**Figure 2.**
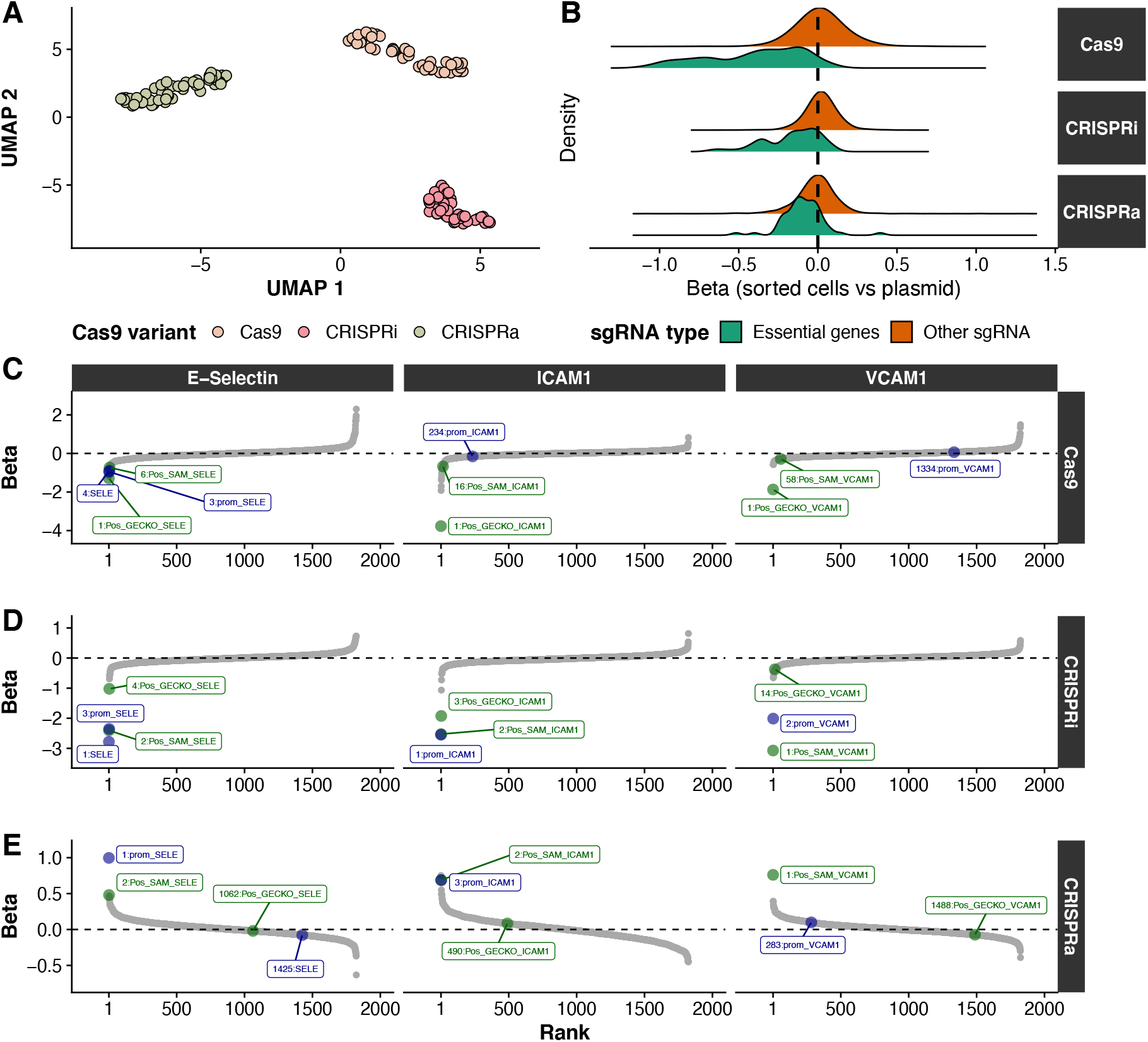
Quality-controls of the pooled CRISPR screens for vascular endothelial cell phenotypes. (**A**) Two-dimensional uniform manifold approximation and projection (UMAP) representation of 148 fluorescence-activated cell sorting (FACS) samples based on the normalized read counts of the top 10% most variable sgRNAs across all samples. (**B**) Density distributions of effect sizes (Beta, *x*-axis) across all Cas9 variants for essential genes and the rest of the sgRNA library. Positive betas indicate that sgRNA are enriched in the cell fractions when compared to the input library, while negative betas indicate a depletion of sgRNA across all samples. We observed a depletion of sgRNA targeting essential genes with all three Cas9 variants. (**C-E**) Rank of all control sgRNAs and targeted CAD variants in the (**C**) Cas9, (**D**) CRISPRi and (**E**) CRISPRa screens for three adhesion proteins: E-selectin (left), VCAM1 (middle) and ICAM1 (right). For each panel, the *y*-axis corresponds to the effect sizes (Beta, comparing top *vs* bottom FACS 10% fractions). For the Cas9 and CRISPRi experiments, we found an enrichment of sgRNAs targeting the coding and promoter sequences of genes encoding adhesion proteins in the bottom 10% cell fractions (negative Betas). In contrast, sgRNAs targeting the promoter of these genes were enriched in the top 10% cell fractions in the CRISPRa experiments. In green and blue, we highlight sgRNAs targeting coding exons and promoters, respectively. The number in front of the name of each control sgRNA indicates its rank in the corresponding analysis.

### Effects of CRISPR knockout, inhibition and activation in teloHAEC

To assess Cas9 efficiency in our experiments, we included in the library 330 sgRNAs against the coding sequence of genes essential for cell viability. For Cas9 and CRISPRi, we found a strong depletion of sgRNAs targeting essential genes among the sequenced FACS cell fractions (Kolmogorov-Smirnov (KS) test *P*<2.2×10^−16^ and *P*=1.6×10^−13^, respectively) (**Figure 2B**). We also noted a minor but significant shift toward depletion in the sgRNA count distribution of essential genes for the CRISPRa experiments (KS test *P*=3.7×10^−6^), potentially due to steric hindrance effects by the dead Cas9 moiety near the transcriptional start site of these genes or the toxic impact of gene over-expression (**Figure 2B**).

As an additional quality-control step in our experiment, we designed sgRNAs against the coding and promoter sequences of *SELE, ICAM1* and *VCAM1,* which encode the three adhesion proteins measured in our FACS assays (**Figure 1F**). We observed significant depletion of sgRNAs targeting coding exons and promoter regions of these genes in the top vs. bottom 10% FACS fractions with Cas9 or CRISPRi (**Figures 2C-D**, **Supplementary Figure 5**). In the CRISPRa experiments, the same sgRNAs were enriched in FACS fractions with high E-selectin, ICAM1 or VCAM1 levels (**Figure 2E**, **Supplementary Figure 5**). The three other endothelial phenotypes measured in our experiments - NO and ROS production, and Ca^2+^ signalling - are physiological readouts that are not the product of a single gene. In the absence of confirmed positive control genes that we could target to validate our system, we carefully calibrated the flow cytometry assays for these readouts using appropriate agonists/inducers (**Methods**). Our screens are sufficiently sensitive to detect sgRNAs targeting CAD loci that have strong effects on these hallmarks of endothelial dysfunction.

Our pooled CRISPR screens identified 51 significant variant-endothelial phenotype results (false discovery rate (FDR) ≤10%) involving 42 different variants located within 26 CAD loci (**Figure 3A** and **Supplementary Table 4**). We found significant results for almost all combinations of Cas9 modality and FACS phenotypes, and most of these results were specific to a single combination (**Figure 3A**). This highlights the importance to test several cellular phenotypes and Cas9 modalities to carry out comprehensive perturbation screens in order to characterize GWAS loci. For 15 CAD loci where we could target all LD proxies with sgRNAs (**Figure 1D** and **Supplementary Table 3**), we detected no significant signals in our CRISPR assays, suggesting that genes within these genomic regions modulate CAD risk through different functions or cell types. When compared with genomic loci with no significant results, CAD loci with at least one significant variant in our CRISPR screens were not better captured by designed sgRNAs (median coverage 73% vs. 78% of LD proxies, Wilcoxon’s test *P*=0.70) but had significantly more LD proxies (median 41 vs. 9 variants, Wilcoxon’s test *P*=1.1×10^−4^).

**Figure 3.**
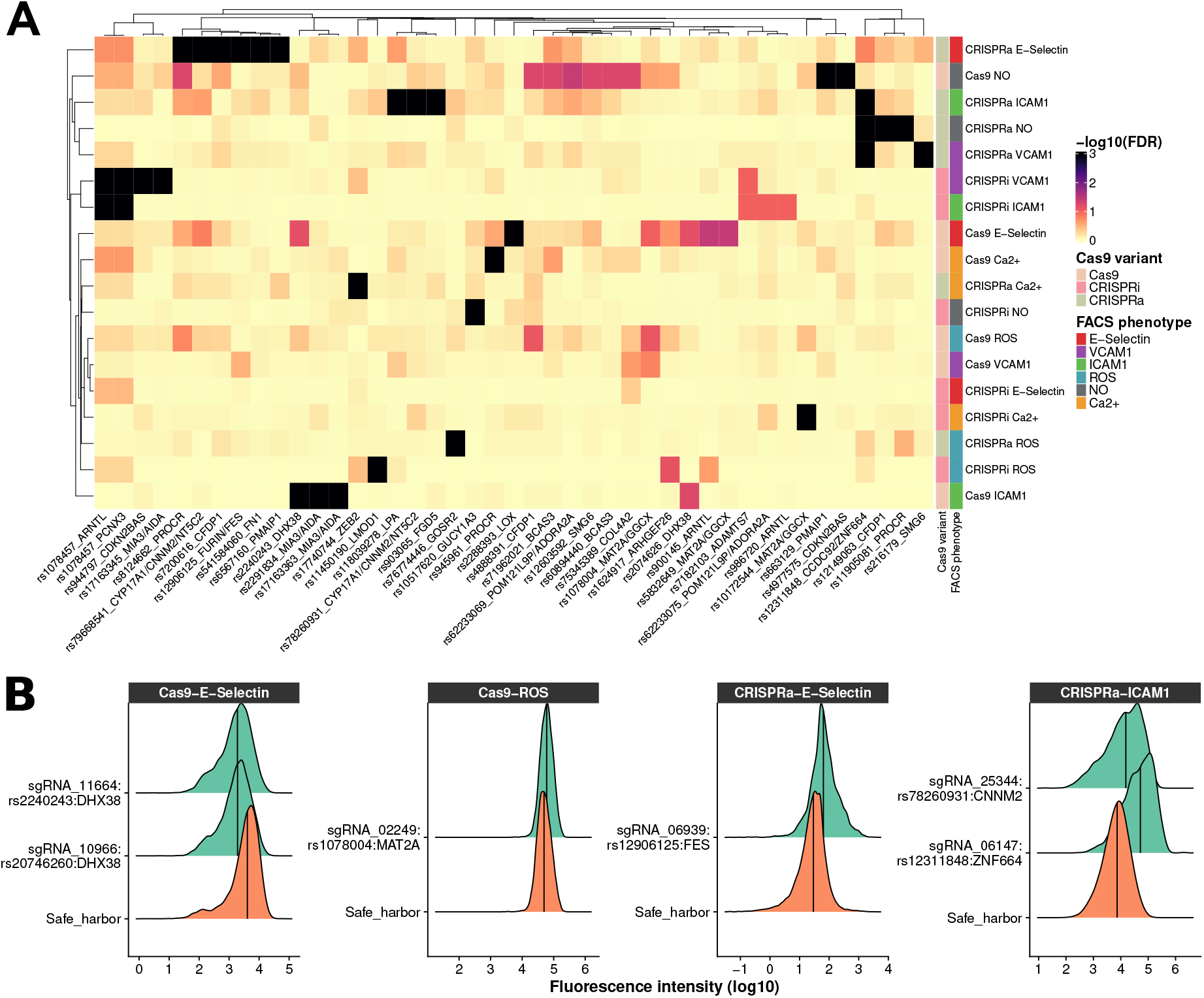
Discovery and validation of CRISPR perturbations that induce atheroprone vascular endothelial cell phenotypes. (**A**) Heatmap of CAD-associated variants that are significant (false discovery rate (FDR) ≤10%) for at least one of six endothelial phenotypes tested in the teloHAEC pooled CRISPR screens. Each row corresponds to a combination of Cas9 variant and cellular readout, and each column corresponds to a CAD variant. For each variant, we added the name of a nearby gene to simplify locus identification, although we do not imply that these genes are causal. Dendrograms of rows and columns represent hierarchical clustering based on euclidean distance. The FDR is capped at 0.1%. (**B**) Validation by flow cytometry of six hits from the pooled CRISPR screens. For each validation, we used the top sgRNA from the pooled CRISPR screens to target the variant/locus with the corresponding Cas9 variant. We compared the distribution of the fluorescence intensity of the cellular markers (*x*-axis) between the sgRNA identified in the screens and a safe harbor negative control sgRNA. We assessed statistical significance using the Kolmogorov-Smirnov (KS) test, all validations shown are significant (KS P-value <2.2×10^−16^). Validations were performed in at least three independent experiments for each sgRNA (**Supplementary Table 5**).

Several of the CAD loci identified by GWAS have been implicated in blood lipid metabolism (*e.g. LDLR, APOE, PCSK9*). Because genetic variation within these loci are likely to influence risk through an effect on lipid levels, we did not anticipate identifying them in our endothelial cell functions CRISPR screens. Of the variants that mapped to 10 lipid loci included in our screens, all were negative across the different endothelial phenotypes tested except rs118039278 located in an intron of *LPA* (CRISPRa for ICAM1, FDR<0.001, **Figure 3A**). Although *LPA* is not expressed in teloHAEC, CRISPRa could induce its ectopic expression and the encoded Lp(a) lipoprotein has been shown to induce endothelial dysfunction^22^.

### Validation of a CAD-associated regulatory variant at the *FURIN/FES* locus

To validate our results, we selected eight SNPs at seven CAD loci and performed individual sgRNA infection and FACS experiments (**Supplementary Table 5**). For this validation step, we prioritized variants that were significant for >1 cellular phenotype and that had strong effect sizes in the CRISPR screens. For each experiment, we compared the distribution of the FACS-based cellular phenotype between control sgRNAs and the best sgRNA targeting each selected CAD variant (**Figure 3B**). Across three independent biological replicates, we could validate six of the eight selected SNPs (one-tailed *t*-test *P*<0.05, **Supplementary Table 5**).

To investigate the global transcriptional consequences of genome editing perturbations and elucidate the underlying impact of these loci on endothelial cell biology, we performed RNA-seq for five sgRNAs that passed primary validation. At the *FURIN/FES* locus, the top sgRNA (sgRNA_06939) maps to rs12906125, a variant in strong LD with the CAD sentinel variant rs2521501 (*r*^2^=0.91). rs12906125 is located in the *FES* promoter and overlaps an ATAC-seq peak as well as a H3K27ac-defined enhancer that physically interacts with the *FURIN* promoter (**Figure 4A**)^9^. The same SNP is an eQTL for *FES* in human primary aortic endothelial cells^23^ and arterial tissues from GTEx. In the CRISPRa experiments, we found a significant up-regulation of both *FES* (log2(fold-change (FC))=3.75, adjusted *P*=8.5×10^−173^) and *FURIN* (log2FC=0.78, adjusted *P*=1.5×10^−10^) (**Figure 4B**).

**Figure 4.**
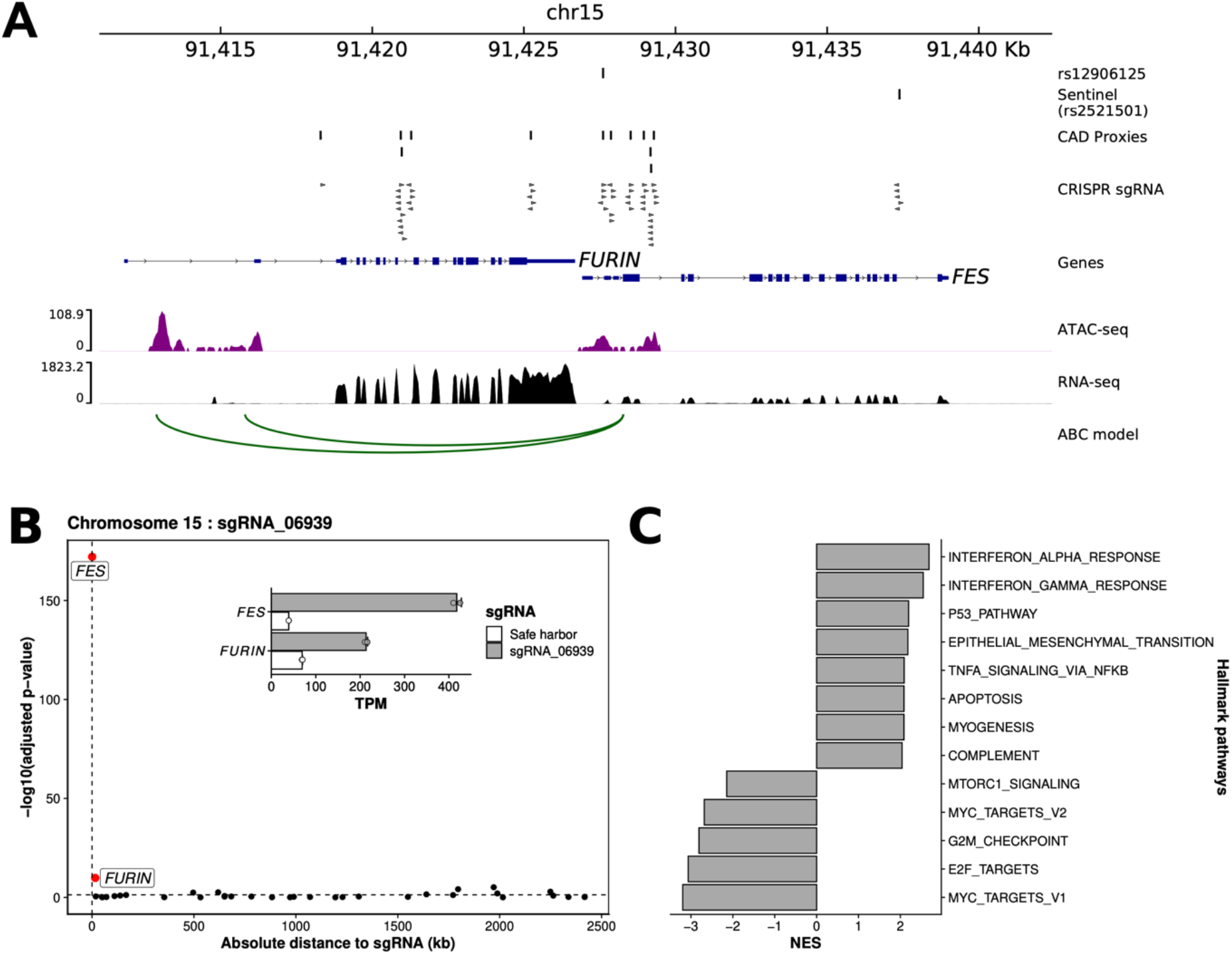
Characterization of a CAD-associated regulatory variant located within an enhancer at the *FURIN/FES* locus. (**A**) CRISPRa perturbations highlighted rs12906125 as a potential regulatory variant for *FURIN* and *FES*. The variant overlaps an ATAC-seq peak in the promoter of *FES* and a H3K27ac-defined enhancer that physically interacts with the *FURIN* promoter through chromosomal loops predicted by the ABC model applied to teloHAEC Hi-C data^9,64^. (**B**) Within a 2.5-Mb window, *FES* and *FURIN* are the top two differentially expressed genes when targeting rs12906125 by CRISPRa in teloHAEC. The inset plot shows the induction of both *FES* and *FURIN* expression with sgRNA_06939 when compared to the control safe harbor sgRNA. (**C**) Enrichment results of differentially expressed genes for the MSigDB (H) hallmark gene-sets, comparing RNA-seq data for CRISPRa experiments with sgRNA_06939 targeting variant rs12906125 to a control safe harbor sgRNA. Only pathways with a Benjamini-Hochberg-corrected P-value <0.05 and a normalized enrichment score (NES) <-2 or >2 are highlighted. All results from the pathway analyses are presented in **Supplementary Table 6**.

Recently, a different CAD SNP at the same locus, rs17514846, was shown to be an eQTL for *FURIN* in vascular endothelial cells^24^. This variant was not tested in our CRISPR screens because of weaker LD with the sentinel variant rs2521501 (*r*^2^=0.47). *FURIN*, which encodes a proprotein convertase, represents a strong candidate CAD causal gene at this locus: its specific knockdown in human endothelial cells reduced atheroprone characteristics such as monocyte-endothelial adhesion and transmigration^24^. Consistent with these previous *FURIN*-related results, gene-set enrichment analysis (GSEA) of the *FURIN/FES* RNA-seq data highlighted genes implicated in inflammatory responses (interferon, TNFα/TGFβ) and cell cycle regulation (p53, apoptosis) (**Figure 4C** and **Supplementary Table 6**). Although our results are consistent with *FURIN* representing a strong CAD candidate gene, we cannot rule out a role for *FES* given the strong CRISPRa effect (**Figure 4B**). *FES*, which encodes a tyrosine protein kinase that can control cell growth, differentiation and adhesion, has not been implicated in vascular endothelial cell biology.

### Loss of *DHX38* function induces vascular endothelial cell senescence

Two of the validated sgRNAs target synonymous variants in *DHX38* (rs2074626, rs2240243) and mediate Cas9 nuclease effects on E-selectin (**Figures 3A-B**) and VCAM1 (as validated by subsequent analyses, **Supplementary Table 5**). We confirmed the E-selectin result using Cas9 ribonucleoprotein complexes (**Supplementary Figure 6**). In our screens, we also tested but found no significant effects for two *DHX38* missense variants (rs1050361, rs1050362). However, in contrast to the *DHX38* synonymous variants, these missense variants are located in early exons (exons 2-3, with transcripts potentially escaping nonsense-mediated mRNA decay) that are expressed at low levels. *DHX38* encodes a RNA helicase involved in splicing (**Figure 5A**).

**Figure 5.**
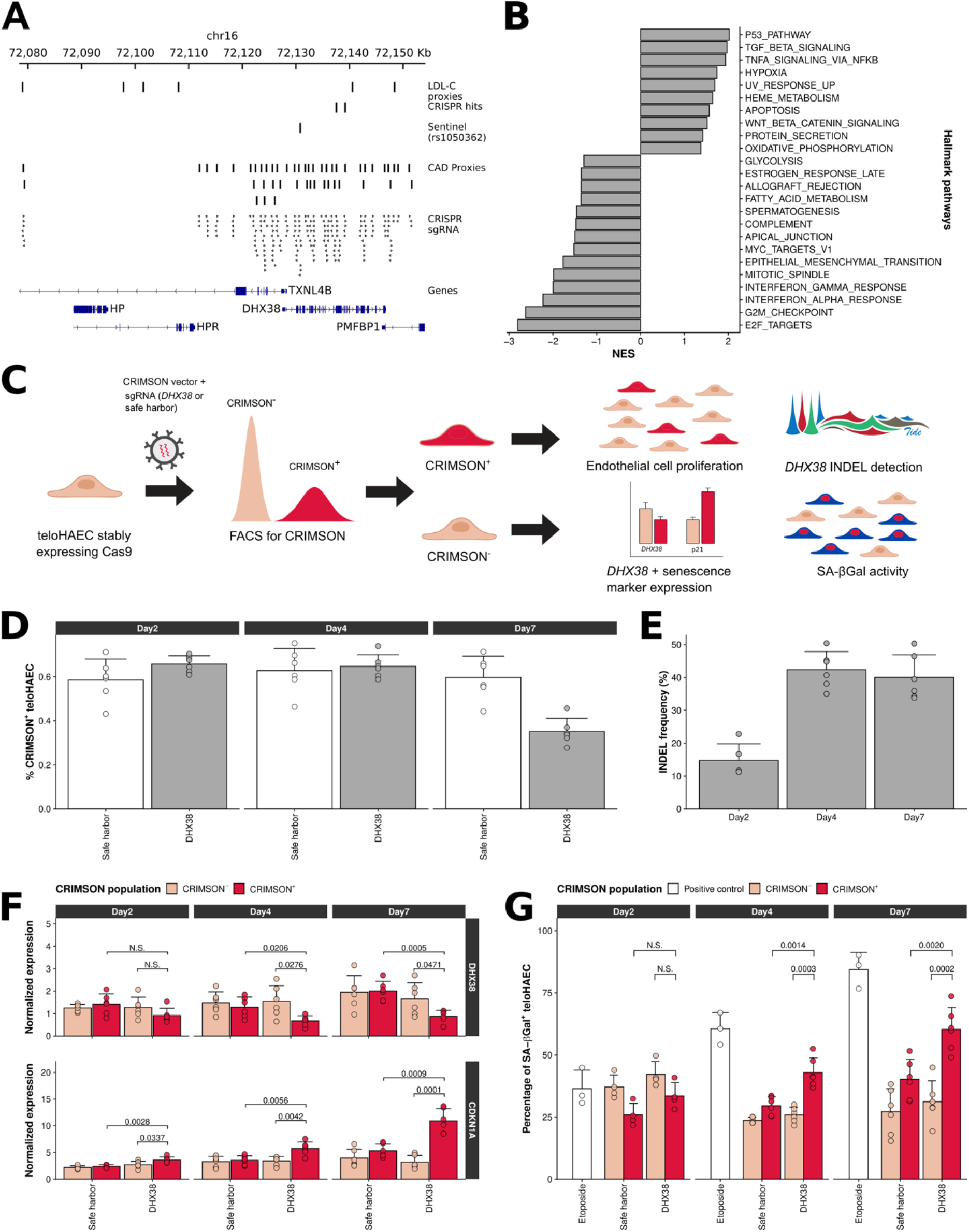
Disruption of *DHX38* induces vascular endothelial cell senescence. (**A**) Perturbations with the Cas9 nuclease highlighted two synonymous variants (rs2074626, rs2240243) in the *DHX38* gene for several endothelial phenotypes. *DHX38* is located downstream of the *HP* and *HPR* genes, which have previously been associated with LDL-C levels. However, the CAD and LDL-C GWAS signals are distinct. (**B**) Gene-set enrichment analysis results for differentially expressed genes identified by RNA-seq in teloHAEC between a sgRNA targeting a *DHX38* coding exon and a safe harbor negative control sgRNA. Only pathways with a Benjamini-Hochberg-corrected P-value <0.05 and normalized enrichment scores (NES) <−1 or >1 are shown. (**C**) Experimental design for the characterization of *DHX38* using the fluorescent marker CRIMSON in place of an antibiotic resistance gene. We did all experiments in teloHAEC that stably express Cas9. We monitored the impact of a *DHX38* sgRNA on cell proliferation, indel induction, gene expression and senescence-associated β-galactosidase (SA-βGal) activity. (**D**) Comparison of endothelial cell proliferation between teloHAEC with a *DHX38* sgRNA or a safe harbor negative control sgRNA. The differences in the number of CRIMSON^+^ cells were not significant two or four days post-infection. However, there were 27% less CRIMSON^+^ cells with *DHX38* sgRNA relative to the safe harbor control at seven days post-infection (Student’s *t*-test *P*-value = 7.3×10^−8^). Results are mean ± standard deviation for 6 replicates for safe harbor and three replicates for two DHX38 targeting sgRNA. (**E**) Quantification of *DHX38* indels by tracking of indel by decomposition (TIDE) analysis. As expected, we found no indels in the CRIMSON^−^ cells (**Supplementary Table 7**). However, in CRIMSON^+^ cells that received a *DHX38* sgRNA, we found indels with an average frequency of 15%, 42% and 40% at day 2, 4 and 7, respectively. Results are mean ± standard deviation for 6 replicates for safe harbor and three replicates for two DHX38 targeting sgRNA. (**F**) Expression levels of *DHX38* and *CDKN1A* in CRIMSON^−^ and CRIMSON^+^ teloHAEC that have received a sgRNA that targets *DHX38* or a safe harbor region (negative control). There were no significant differences in *DHX38* expression levels at day 2. However, at day 4 and 7, *DHX38* was significantly down-regulated and *CDKN1A* was significantly up-regulated in CRIMSON^+^ cells that received the *DHX38* sgRNA. N.S., not significant. We provide Student’s *t*-test P-values when *P*<0.05. Bars are mean normalized expression and error bars represent one standard deviation. (**G**) Quantification of senescent teloHAEC by flow cytometry using senescence-associated β-galactosidase (SA-βGal) staining. At day 4 and 7 post-infection, there were significantly more senescent cells in the CRIMSON^+^ *DHX38* sgRNA experiment than in the CRIMSON’ cells or in the CRIMSON^+^ cells that received the safe harbor sgRNA. We used the DNA damaging agent etoposide as a positive control to induce senescence. N.S., not significant. We provide Student’s *t*-test P-values when *P*<0.05. Results are mean percentage SA-βGal^+^ teloHAEC and error bars represent one standard deviation.

In the RNA-seq experiments with a sgRNA targeting *DHX38* (seven days post-infection, TNFα treatment), the gene was not down-regulated and we found few reads with Cas9-mediated indels (<2%), yet a strong gene expression signature suggesting an effect on cell proliferation with the modulation of genes involved in the p53, G2/M checkpoint and E2F target genes pathways (**Figure 5B**). Together, these results suggest a complex scenario: two sgRNAs targeting Cas9 nuclease to *DHX38* exons produce robust cell adhesion phenotypes as well as a cell cycle-related gene expression signature without apparently introducing many indels nor impacting *DHX38* expression levels. To reconcile these observations, we hypothesized that endothelial cells with *DHX38* detrimental indels have a growth disadvantage and induce a response in surrounding cells without *DHX38* indels through paracrine signalling. To test this model, we replaced the antibiotic resistance marker by a fluorescence protein (CRIMSON) in the sgRNA vector so that we can sort and characterize at different timepoints teloHAEC stably expressing Cas9 that have or not received a *DHX38* sgRNA (**Figure 5C**). While the fraction of CRIMSON^+^ cells is similar for safe harbor and *DHX38* sgRNAs two- and four-days post-infection, it is significantly lower after seven days (**Figure 5D**). This observation is aligned with our pooled CRISPR screens results since cells were also tested for endothelial dysfunction phenotypes seven days post-infection. Although we did not capture many *DHX38* indels in the RNA-seq experiment, we could detect a high frequency of indels (15-40%) in CRIMSON^+^ cells already two days post-infection (**Figure 5E** and **Supplementary Table 7**). Importantly, we also measured a down-regulation of *DHX38* expression levels in CRIMSON^+^ cells, confirming that *DHX38* is likely the gene at this CAD locus that mediates the endothelial phenotypes (**Figure 5F**).

Our hypothesis would be consistent with vascular endothelial cell senescence and the senescence-associated secretory phenotype (SASP)^25,26^. In CRIMSON^+^ cells with *DHX38* sgRNA, we measured an up-regulation of *CDKN1A* (encoding the CDK2 inhibitor p21^WAF1/Cip1^) and detected a higher number of cells with β-galactosidase activity when compared to CRIMSON^−^ cells or CRIMSON^+^ cells with a safe harbor sgRNA (**Figures 5F-G**). These characteristics are hallmarks of cell senescence. Activation of the senescence program is specific to *DHX38* and not a general response to DNA damage induced by this particular sgRNA as four different sgRNAs targeting two different *DHX38* exons impaired endothelial functions in the CRISPR screens. Collectively, our observations point towards the induction of senescence in teloHAEC with dysregulated *DHX38* expression and help explain the results from our RNA-seq experiment. In the cell population profiled by RNA-seq, we found few *DHX38* indels and did not detect *DHX38* down-regulation because cells with *DHX38* detrimental indels underwent senescence-mediated growth arrest. However, these *DHX38*-edited senescent cells secreted pro-inflammatory molecules as part of the SASP which, in combination with the TNFα added to the cell medium, activated a transcriptional response in teloHAEC without *DHX38* edits.

### A transcriptional signature of senescence triggered by CRISPRa perturbations at three CAD loci

To assess the role of vascular endothelial cell senescence in explaining our CRISPR screens results, we re-analyzed our RNA-seq data using senescence and SASP curated gene sets^27^. This analysis did not yield significant enrichments for sgRNAs that target *DHX38* or *MAT2A* (**Figure 6A**). Despite our results that implicate *DHX38* in senescence (**Figure 5**, CRIMSON experiments without TNFα), we did not expect to find a gene expression signature of senescence in the RNA-seq experiment (with TNFα) because too few cells had *DHX38* indels (see above). Additionally, the SASP was masked by the presence of exogenous pro-inflammatory TNFα, which we added to the cell medium to validate the E-selectin phenotype (**Figure 3B**). For *MAT2A,* targeting Cas9 to the synonymous variant rs1078004 increased ROS production in TNFα-treated teloHAEC (**Figure 3C**). *MAT2A* encodes a methionine adenosyltransferase that is responsible for the biosynthesis of S-adenosylmethionine, a precursor of the potent antioxidant glutathione^28^. The analysis of the *MAT2A* RNA-seq experiment (with TNFα) revealed a profile very similar to *DHX38:* we found few *MAT2A* indels, the gene was not differentially expressed, and we did not detect a signature of senescence (**Figure 6A**). Thus, it is possible that *MAT2A* edits decrease *MAT2A* expression, increased oxidative stress and trigger senescence^25,26^, but that we would only observe this response by enriching for cells with *MAT2A* sgRNA in medium without exogenous TNFα, as we did for *DHX38* sgRNA (**Figure 5**).

**Figure 6.**
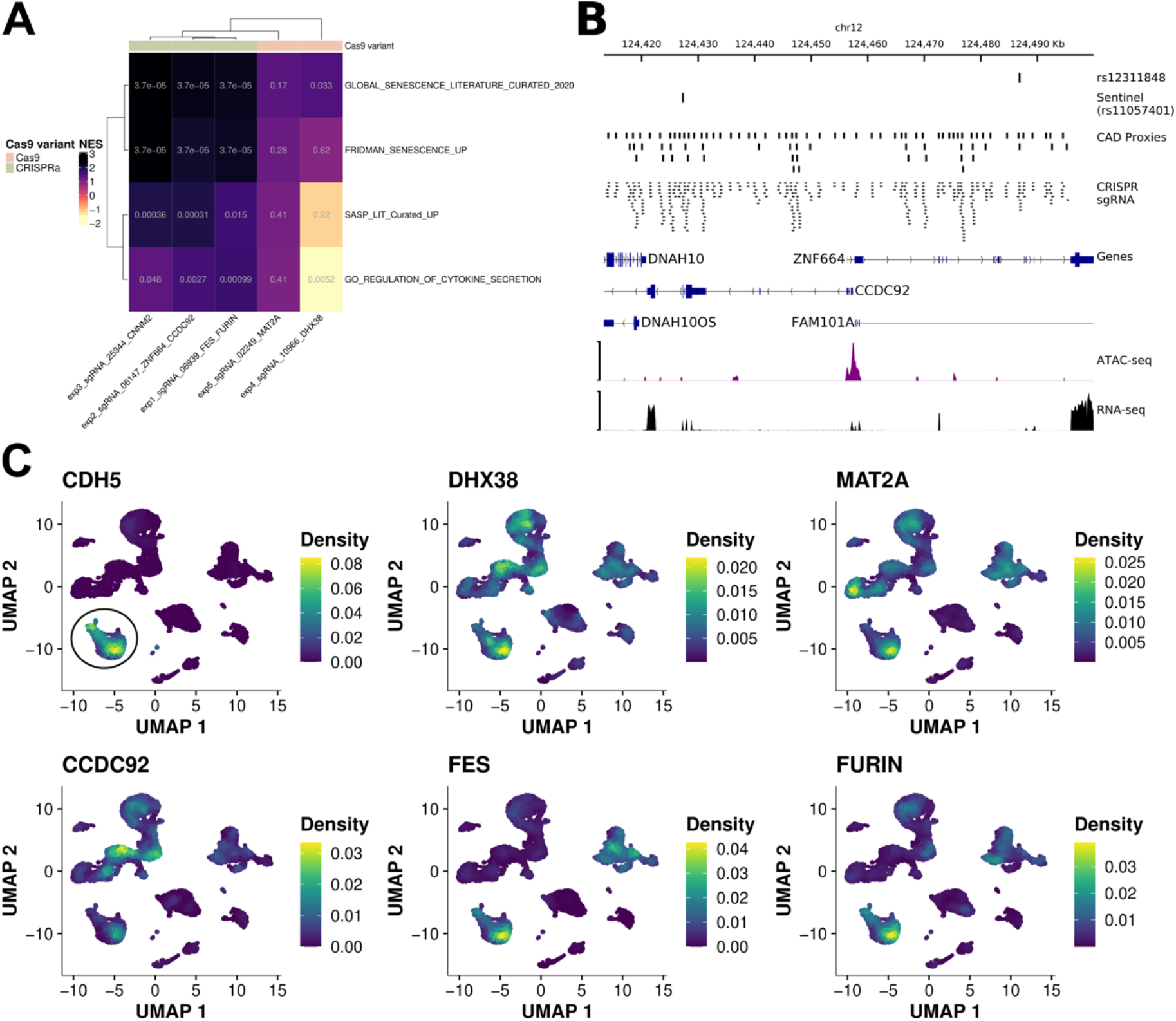
CRISPRa at the *FURIN/FES, CCDC92/ZNF664* and *CNNM2* loci activates a gene expression signature of senescence. (**A**) Gene-set enrichment analyses of the five RNA-seq experiments for four curated gene sets that capture senescence or the SASP ^27^. The three CRISPRa experiments are highly significant for the senescence pathways. In each cell of the heatmap, the color and number indicate, respectively, the normalized enrichment score (NES) and the Benjamini-Hochberg adjusted P-value. (**B**) Locus view for the CAD locus with nearby genes *ZNF664* and *CCDC92.* We provide the position of the sentinel CAD variant (rs11057401) and the functional variant identified in the pooled CRISPR screen (rs12311848). The LD proxies and sgRNAs tested are also shown. ATAC-seq and RNA-seq data in resting teloHAEC are from ref. ^9^. *DNAH10, DNAH10OS, FAM101A,* and *ZNF664* were not differentially expressed in the CRISPRa experiment. (**C**) Uniform manifold approximation projection (UMAP) for 11,756 cells from human right coronary arteries analyzed by single-cell RNA-sequencing^65^. We color-coded cells based on the level of expression of candidate causal CAD genes identified and characterized in this study. We used the expression of the endothelial cell marker gene *CDH5* (encoding VE-Cadherin) to identify endothelial cells (circle in top left panel). All five candidate genes are expressed in human vascular endothelial cells from coronary arteries.

Analysis of the RNA-seq data for the three CRISPRa experiments showed a strong senescence signature (**Figure 6A**). While it has recently been reported that CRISPRa can lead to non-specific transcriptional effects such as the up-regulation of *IL6*^29^, we used safe harbor sgRNAs to control for such effects and *IL6* was not differentially expressed in our experiments. Additional controls suggested specificity of our CRISPRa results (**Supplementary Figure 7**). Beside the *FURIN/FES* locus described above, the two other CRISPRa experiments targeted intronic variants in *ZNF664* (rs12311848) and *CNNM2* (rs78260931). Across these three CAD loci, the CRISPRa experiments shared 734 differentially expressed genes and highlighted biological pathways that are relevant to senescence, such as the down-regulation of cell cycle-related genes (**Supplementary Tables 6** and **8**).

Targeting CRISPRa at rs12311848 did not increase the expression of *ZNF664* but the expression of *CCDC92,* a gene located 82 kb upstream (log2FC=0.74, adjusted *P*=9.2×10^−5^, **Figure 6B**). The sentinel CAD variant identified by GWAS at this locus is rs11057401, a missense variant in *CCDC92.* We targeted four sgRNAs at rs11057401 but did not detect significant effects in the CRISPR screens. This result suggests that CRISPRa gain-of-function experiments are necessary to detect the impact of *CCDC92* on endothelial dysfunction and senescence. *CCDC92* has been found to be over-expressed in senescent cells^30^; accordingly, we detected increased *CCDC92* expression in the *FURIN/FES* (sgRNA_06939, log2FC=0.6, adjusted *P*=3.3×10^−5^) and *CNNM2* (sgRNA_25344, log2FC=1.9, adjusted *P*=1.8×10^−42^) CRISPRa experiments. *CCDC92* over-expression is specific to these sgRNAs as a CRISPRa experiment with a “silent” sgRNA targeting the *LPL* locus did not induce its expression (**Supplementary Figure 7B**). Finally, although we measured a robust transcriptional signature of senescence when targeting CRISPRa at the *CNNM2*-rs78260931 locus (**Figure 6A**), we found no evidence of differential expression for nearby genes (in *cis,* the closest differentially expressed gene was *NFKB2* located 568 kb away (log2FC=0.33, adjusted *P*=0.008)). We also manually inspected the sequence reads that mapped to the *CNNM2* region but did not find un-annotated genes that were differentially expressed. Thus, based on our results, we cannot prioritize a candidate causal gene at this CAD locus, but emphasize that the effect is specific to this region and not a sgRNA-specifc artifact because three of the four sgRNAs that we targeted at rs78260931 gave consistent results in the CRISPRa-ICAM1 screen.

## DISCUSSION

As for most complex human diseases, many GWAS loci associated with CAD do not include obvious candidate causal genes nor implicate known pathophysiological mechanisms. To elucidate their mechanisms and gain insights into atherosclerosis, we carried out multiple CRISPR screens to test if CAD variants impact vascular endothelial functions. By combining six different endothelial cell readouts and three Cas9 modalities, we identified 42 variants at 26 CAD loci (**Figure 3A**). This list is depleted of variants that modulate CAD risk through an effect on lipid metabolism and enriched for loci of unknown functions (**Supplementary Table 3**). We found variants near *ARHGEF26* and *ADAMTS7,* genes previously implicated in leukocyte transendothelial migration^5^ and endothelial cell angiogenesis^31^, respectively. We also retrieved rs17163363, an intronic variant in *MIA3* that controls the expression of *AIDA* in endothelial cells^9^.

There were also variants and genes that we expected to find but did not recover. For instance, we did not identify rs17114036, a likely functional variant that controls the expression of *PLPP3* in endothelial cells, although this negative result may be because the underlying enhancer requires hemodynamic stress to be active^10^. Furthermore, our screens did not yield variants at CAD loci that include *PECAM1* (adhesion protein CD31) and *NOS3* (endothelial NO synthase), two genes with important roles in endothelial cells. As for *PLPP3*, it might be that we did not activate endothelial cells with the right stimulus to detect the functional impact of these variants/genes in our assays. It is also possible that some loci will require the precise engineering of alleles (*e.g.* using base editing) to detect a cellular phenotype, or that the phenotypic effect of a variant at the cellular level is too low to distinguish a true signal from the experimental noise inherent to any large-scale omics approach.

We designed our sgRNA library using a variant-focused approach. Our rationale was that the identification of causal variants by CRISPR perturbations would lead us to the causal genes and biological pathways. This strategy worked at a few CAD loci (*e.g. MIA3/AIDA*^9^). At the *FURIN/FES* locus, we found a candidate regulatory variant that was also prioritized using orthogonal methodologies (**Figure 4**)^23^. However, it is also likely that some of the findings from our CRISPR screens result from loss- or gain-of-function effects on causal genes independently of the causal variants. For instance, we identified and validated sgRNAs near synonymous variants in *DHX38* and *MAT2A* using the Cas9 nuclease. While synonymous variants can have phenotypic consequences, it is more likely that these variants are in LD with the causal variants but were captured in our screens because they targeted loss-of-function indels to the *DHX38* and *MAT2A* coding sequences. Similarly, ectopic activation or inhibition of gene expression by CRISPRa and CRISPRi can highlight potential nearby causal genes (*e.g. CCDC92)* even if the sgRNAs do not directly overlap causal variants.

Vascular endothelial cell senescence emerged as an important response in our CRISPR perturbation screens. Senescent endothelial cells are characterized by growth arrest, but also a pro-inflammatory and atheroprone phenotype that involves increased production of adhesion molecules and ROS, and reduced NO bioavailability^32^. For *DHX38* with Cas9, we had to design a TNFα-free CRIMSON-based strategy to confirm senescence (**Figure 5**). Conversely, the RNA-seq results for the CRISPRa experiments (without TNFα) at *FURIN/FES, CCDC92/ZNF664* and *CNNM2* were less ambiguous, with a clear gene expression signature indicating senescence (**Figure 6A**). Our controls do not support the hypothesis that this difference is due to non-specific effects of Cas9 or CRISPRa (**Supplementary Figure 7**). Instead, the presence of TNFα, which can induce premature endothelial cell senescence^33^, has led to a stronger growth arrest phenotype in the presence of Cas9-mediated indels in *DHX38* (and maybe also *MAT2A*).

While *FURIN* has been shown to influence endothelial functions^34^, most other genes (*FES*, *DHX38, MAT2A,* and *CCDC92)* have been less studied. Supportive of their role *in vivo,* we re-analyzed single-cell RNA-seq data from human atherosclerotic right coronary arteries and confirm that these genes are expressed in human vascular endothelial cells (**Figure 6C**). Focusing specifically on *CCDC92,* the same CAD variant is also associated with waist-to-hip ratio, and knockdown of *CCDC92* expression in immortalized mouse OP9-K stromal cells impaired adipogenesis^35^. This result prompted the authors to propose a role for *CCDC92* in adipocyte differentiation and insulin secretion, although the authors did not exclude senescence as a possible mechanism. In a different study, *CCDC92* over-expression reduced Ebola viral infections in HEK293 and endothelial HUVEC by blocking the virus without killing the host cells^36^. It was not investigated, but senescence has been proposed as an antiviral strategy^37^. Given that *CCDC92* was up-regulated both in *cis* and *trans* in our CRISPRa experiments, it suggests that it is a key gene located in a CAD GWAS locus that can impact normal endothelial functions.

Endothelial cell senescence is both a physiological and pathological process^38^. In health, it signals the system for vascular endothelium repair. Senescence also increases with age and in response to traditional CAD risk factors. When it overcomes the regeneration capacity of the system or upon stress, senescence causes endothelial dysfunction and can lead to vascular diseases. Senescent cells accumulate at the sites of atherosclerosis in human blood vessels^39,40^ and their selective elimination using transgenic strategies or drugs (senolytics) delays atherogenesis progression in mice^41^. Our data suggest that a subset of variants associated with CAD in humans affect key endothelial functions, potentially by inducing premature senescence. This observation links a large body of literature that has implicated senescence in atherosclerosis with a genetic program that modulates endothelial functions. As clinical trials to test the efficacy of senolytics on vascular diseases are now in discussion^42^, it will be important to explore whether specific CAD variants or polygenic scores are predictive of their clinical response.

## ONLINE METHODS

### Design of the sgRNA library

We retrieved 92 sentinel genetic variants associated with coronary artery disease (CAD) at genome-wide significant levels (P-value ≤5×10^−8^) from four GWAS meta-analyses available at the time of the design of this experiment^5,6,18,19^. For the design of the sgRNA library, we included all sentinel variants as well as variants in strong LD (*r*^2^ >0.8 in the 1000 Genomes Project European-ancestry populations). For each variant - sentinel and LD proxy - we identified all possible sgRNA in a 100-bp window centered on the variant itself. We prioritized sgRNA with the highest predicted quality using the CRISPR OffTarget Tool (version 2.0.3)^43^ with a Targeting_guide_score ≥ 20 and the “matches with 0 mismatches” = 1 and “matches with 1 mismatch” = 0 settings. We discarded sgRNA that overlapped heterozygous variants, indels and/or multi-allelic variants in the teloHAEC genome (build hg19). We selected sgRNA targeting essential genes from a previously published study^44^. For potential positive control genes (*SELE, SELP, ICAM1, VCAM1, PECAM1, NOS3, VWF, SOD2, SOD3, GPX3, CAT, ITPR1, ITPR2, ITPR3, ATP2A2, ATP2A3, PLN, CAV1,* and *TRPV4),* we selected sgRNA from the Human GeCKOv2 CRISPR knockout pooled library^45^. We also selected sgRNA that targeted the promoter (300-bp window before the transcriptional start site) of positive control genes for the CRISPRa (dCas9-VP64) experiments. For all selected loci (variants, coding sequences, gene promoters), we retained the five top scoring sgRNA for the library design. Finally, we added two sgRNA for the *SELE* locus (SELE_g1, SELE_g2) that we frequently use to validate TNFα stimulation. This resulted in a final library of 8051 sgRNA (**Supplementary Table 2**).

The sgRNA were synthesized in duplicates by Agilent Technologies (Cat-#: G7555B) to accommodate the specific requirements of the Cas9/dCas9-KRAB and dCas9-VP64 (specific MS2 tracrRNA) experiments. We amplified each specific pool of oligonucleotides as previously described^16^, with the following small modifications: we performed two PCR using NebNext High fidelity Master mix (Cat-#: M0541L). The first PCR was used to amplify each pool separately using 2.5ng of pooled oligonucleotides and 500nM of each primer (for the Cas9/dCas9-KRAB library, we used U6_subpool_fwd and Guide_CM_barcode1_rev; for the dCas9-VP64 library, we used U6_subpool_fwd and Guide_MS2_Barcode2_rev). Cycling conditions for PCR1 were 98°C for 30 sec, then 15 cycles of 98°C for 10 sec; 55°C for 10 sec; 72°C for 15 sec and a final step of 72°C for 2 min and a 10°C hold. We performed the second PCR to add homologous sequences, using the U6_screen_fwd and Tracr_rev oligonucleotides for the Cas9/dCas9-KRAB library, and the U6_screen_fwd and Tracr_MS2_rev oligonucleotides for the dCas9-VP64 library, in both cases using ⅕ of PCR1 as template. Cycling conditions for PCR2 were 98°C for 30 sec, then 10 cycles of 98°C for 10 sec; 55°C for 10 sec; 72°C for 15 sec and a final cycle of 72°C for 2 min and 10C hold. See table **Supplementary Table 9** for primer details.

After gel extraction and PCR purification, we performed Gibson assembly in both respective vectors (pHKO9-Neo and lentisgRNA(MS2)-zeo backbone addgene 61427). For pHKO9-Neo, we replaced the Crimson fluorescent gene in the pHKO9-Crimson-CM vector (gift from Dan Bauer’s lab) by a neomycin resistance (NeoR) sequence. Briefly, we amplified the NeoR gene from our pCas9-Neo vector^46^ using BsiWI-Neo-Fwd and MluI-Neo_rev primer (**Supplementary Table 9**). After digestion by BsiWI and MluI, we cloned the segment in pHKO9-Crimson_CM, which had been digested with BsiWI and MluI. We amplified each library using ten independent maxi-preparations (Macherey-Nagel cat# 740424). To control the quality of both libraries, we sequenced them on an Illumina HiSeq4000 instrument and calculated the Gini index, which summarizes read distribution across sgRNA in a given pool. For a good-quality sgRNA library, the expected Gini index is ≤0.2, and we obtained Gini indexes of 0.050 and 0.052 for the Cas9/dCas9-KRAB and dCas9-VP64 library, respectively.

### Engineering of teloHAEC cell lines to stably express Cas9 variants

TeloHAEC are immortalized human aortic endothelial cells obtained by over-expressing telomerase (ATCC CRL-4052). These cells have a normal female karyotype (46;XX) and exhibit many of the properties and functions of human vascular endothelial cells^9^. We generated our teloHAEC cells models expressing either Cas9, dCas9-KRAB or dCas9-VP64 + MPHv2 using Addgene vectors #52962, #46911, #61425 and #89308, and lentiviral infection as previously described^46^.

### Pooled CRISPR screen experiments

We produced four batches of lentiviruses for each sgRNA library pool (Cas9/dCas9-KRAB, dCas9-VP64). We infected each teloHAEC cell line (Cas9, dCas9-KRAB, dCas9-VP64) at a multiplicity of infection of 0.3 using each batch of viruses separately. Following viral infection, we selected cells using zeocin (teloHAEC-dCas9-VP64) or G418 (teloHAEC-Cas9/-dCas9-KRAB) for five (teloHAEC-dCas9-VP64) or seven days (teloHAEC-Cas9/-dCas9-KRAB) in vascular cell basal medium (ATCC PCS-100-030) to remove any cells that did not incorporate a vector. After selection, we stimulated cells expressing Cas9 or dCas9-KRAB using TNFα (10ng/μl) for four hours to induce a pro-inflammatory response; we did not stimulate cells expressing dCas9-VP64, reasoning that the VP64 transcriptional domain should activate gene expression. Following TNFα stimulation, we immunostained cells (around 50M cells) with antibodies linked to phycoerythrin for adhesion molecules (E-selectin (BD BIOSCIENCES Cat-#: 551145), VCAM-1 (Cat-#: 12-1069-42), ICAM-1 (Cat-#: 12-0549-42)) or we incubated with fluorescent dye-based reagents for endothelial signaling markers: (nitric oxide (NO) (DAF-FM Diacetate, Cat-#: D23844)), reactive oxygen species (ROS) (CM-H2DCFDA, Cat-#: C6827), calcium signaling (Fura Red, Cat-#: F3021)). We calibrated the FACS assays with positive control treatments to make sure that we could robustly detect changes in the measured phenotypes. Antibodies and fluorescent dye-based reagents were titrated to use optimal concentrations. We also quantified how teloHAEC were responding to ionomycin for calcium signaling, to sodium nitroprusside for NO and to TNFα for reactive oxygen species. For adhesion molecules, we utilized sgRNA targeting coding exons and promoter regions of *SELE, ICAM1* and *VCAM1* as positive controls. Unless otherwise stated, we purchased all antibodies and dyes from ThermoFisher Scientific. Subsequently, we sorted stained cells by flow cytometry on a BD FACSARIA FUSION flow cytometer to collect the top and bottom 10% of fluorescently labeled cells. FACS traces were generated with FlowJo (BD Biosciences). We extracted genomic DNA from both top and bottom 10% cell fractions separately (around 5M cells in each fraction) using the QIAGEN DNeasy Blood and Tissue kit (Cat No. 69504) according to manufacturer’s instructions.

### Amplification and sequencing of pooled CRISPR experiments

We amplified sgRNA sequences from genomic DNA via PCR, followed by a cleanup step using the QIAGEN QIAquick PCR purification kit (Cat-#: 28104) according to the manufacturer’s instructions. We used the primer sequences and PCR settings as previously described in ref. ^16^. We created sequencing libraries using Illumina TruSeq adapters according to the manufacturer’s protocols. We sequenced the libraries on an Illumina Hiseq4000 instrument at the McGill University and Genome Quebec Innovation Centre (MUGQIC). Generally 6 samples were multiplexed per sequencing lane for a target read coverage of ~500 reads per sgRNA per sample (**Supplementary Figure 4B**).

### Computational analysis of pooled CRISPR screen data

We processed raw sequencing data from the BCL to the FASTQ format using bcl2fastq at MUGQIC. Raw FASTQ reads were quality-controlled using FastQC (https://www.bioinformatics.babraham.ac.uk/projects/fastqc/) and MultiQC^47^. We performed downstream analysis of sgRNA sequencing data using MAGECK (v.0.5.9)^48^. We quantified sgRNA sequences using MAGECK count against the list of sgRNA sequences in the library (**Supplementary Table 3**), allowing for no mismatches in the sgRNA sequence. We then tested the difference in sgRNA counts between cell fractions or conditions using MAGECK maximum likelihood estimation (*mle*) method with median normalization^49^.

### UMAP representation of sample-level count data

We normalized raw sgRNA counts using variance stabilizing transformation (vst) in DESeq2^50^. To account for baseline differences between plasmid preparations, we further normalized samples to their respective vector library by dividing the vst normalized sgRNA count by the vst normalized count of the Cas9/dCas9-KRAB or dCas9/VP64 library, respectively. We calculated principal components using the top 10% most variable sgRNA (805 sgRNA) across all cell sorted samples based on normalized counts. Next, we used the loadings from the first three principal components in Uniform Manifold Approximation and Projection (UMAP)^51^ to create a two-dimensional embedding of the normalized sgRNA count data. Each dot in the UMAP plot represents one sequenced sample (top or bottom 10% of stained cells).

### Analysis of sgRNAs targeting essential genes

To test for potential effects of sgRNA on endothelial cell death and proliferation, we compared sgRNA counts of all samples across the same cellular model (Cas9, dCas9-KRAB, dCas9-VP64) against the respective baseline vector library sgRNA count using MAGECK *mle.* We used sgRNAs targeting essential genes in the teloHAEC Cas9 cellular model as positive controls.

### Single sgRNA validation

We individually cloned each sgRNA for validation as previously described^52^. We produced lentiviruses, infected cells, performed antibiotic selection, and stained cells as for the pooled CRISPR screen. We analyzed cells using flow cytometry (BD FACSCelesta (BD Biosciences, San Jose, CA, USA) equipped with a 20 mW blue laser (488 nm), a 40 mW red laser (640 nm), and a 50 mW violet laser (405 nm). For each experiment, we measured the mean fluorescent intensity (MFI) obtained for sgRNA of interest and compared it with the MFI for control sgRNA (safe-harbor and/or scrambled sgRNA). We performed each experiment at least three times. For statistical analyses, we used Student’s *t*-test and determined that a sgRNA had a significant effect on the measured phenotype when a one-tailed P-value ≤0.05.

For *DHX38* Crimson experiments, we individually cloned each sgRNA in pHKO9-Crimson-CM vector (gift from Dan’s Bauer lab). We produced lentiviruses, infected cells and performed flow cytometry on a BD FACSARIA FUSION flow cytometer at day 2, day 4 and day 7 post-infection. We analyzed the percentage of Crimson positive cells and we sorted Crimson positive and negative cells to extract RNA in each fraction. We extracted total RNA using RNeasy Plus Mini Kit (Qiagen cat #: 74136). We measured RNA integrity and concentration using Agilent RNA 6000 Nano II assays (Agilent Technologies) on an Agilent 2100 Bioanalyzer and Take3 on Cytation V (Biotek). We reverse transcribed 750ng of total RNA using random primers and 1 U of the MultiScribe Reverse Transcriptase (Applied Biosystems) in a 20 μL reaction volume at 100 mM dNTPS and 20 U of RNase inhibitor with these three steps: 10 min at 25 °C, 120 min at 37 °C and 5 min at 85 °C. We followed the MIQE guidelines to assess quality and reproducibility of our qPCR results ^53^. We performed qPCR in triplicates for all samples using: 1.25 μL of cDNA (1/50 dilution), 5 μL of Platinum SYBR Green qPCR SuperMix-UDG (Life Technologies) and 3.75 μL of primer pair mix at 0.8 μM on a CFX384 from Biorad. We used the following thermal profile: 10 min at 95 °C, and 40 cycles of 30 s at 95 °C, 30 s at 55 °C and 45 s at 72 °C. We carried out melting curve analyses after the amplification process to ensure the specificity of the amplified products. We also simultaneously performed qPCR reactions with no template controls for each gene to test the absence of non-specific products. Cq values were determined with the CFX Manager 3.1 (Bio-Rad) software and expression levels were normalized on the expression levels of the house-keeping genes TATA-box binding protein (*TBP*), hypoxanthine-guanine phosphoribosyltransferase (*HPRT*), and glyceraldehyde 3-phosphate dehydrogenase (*GAPDH)* using the ΔΔCt method. The primer sequences are in **Supplementary Table 8**.

### PCR for determination of CRISPR-Cas9-induced indels

We isolated gDNA using QuickExtract DNA Extraction Solution (Epicentre, QE0905) from 1×10^5^ cells. We used 100 or 200 ng of gDNA as a template for PCR reaction with the corresponding primers (see **Supplementary Table 9**). gDNA from parental teloHAEC cells was used as control. Obtained PCR products were analysed by electrophoresis on a 1% agarose gel prior to Sanger sequencing. We used TIDE (tracking of indels deconvolution) software for analysis^54^.

### Assays for assessment of cell senescence

Using the same experimental design (DHX38-Crimson), we performed beta-galactosidase staining using the CellEvent Senescence Green Flow Cytometry Assay Kit from Invitrogen at day 2, day 4 and day 7 following the manufacturer’s protocol. Briefly, we trypsinized and we fixed the cells with 2% paraformaldehyde solution during 10 minutes at room temperature, washed them in 1%BSA/PBS and incubated for 1h30 in 1/500 working solution. After incubation, we washed the cells with 1%BSA/PBS and analyzed them by flow cytometry. We measured β-galactosidase fluorescence signal in positive and negative Crimson cells independently. As positive control, non-infected cells were treated with 20 μM of Etoposide (Sigma, E1383-25) for 2, 4 and 7 days.

### Transcriptome data analysis

For RNA-seq analysis, we extracted RNA using RNeasy plus mini kit from Qiagen (cat #: 74136). RNA-seq experiments were carried out by the Centre d’Expertise et de Services Genome Quebec using rRNA-depleted TruSeq stranded (HMR) libraries (Illumina) on an Illumina Hiseq 4000 instrument (paired-ends, 100-bp reads) and by The Center for applied Genomics (Toronto) using rRNA-deletion library prep on an Illumina NovaSeq-SP flow cell. We quality-controlled raw fastq files with FastQC and multiQC^47^. We used kallisto (v. 0.46.0) to quantify transcript abundances^55^ against ENSEMBL reference transcripts (release 94) followed by tximport to calculate gene-level counts^56^. We utilized regularized log-transformation (rlog) in DESeq2^50^ as input for principal component analysis (PCA). DESeq2^50^ was further used to identify differentially-expressed genes between teloHAEC cell models (Cas9, dCas9-VP64) infected by lentiviruses with safe-harbor sgRNA or sgRNA identified in the pooled CRISPR screens. We excluded genes expressed with less than 10 reads across all samples from the analysis. We performed shrinkage for effect size estimates using apeglm using the lfcShrink method^57^. Genes differentially expressed with a Benjamini-Hochberg adjusted p-value ≤ 0.05 were considered significant. Gene set enrichment analysis was performed using the R package fgsea using 100,000 permutations against the Hallmark gene sets from msigdbr (https://igordot.github.io/msigdbr/)^58,59^. We quantified short indels in the RNA-seq data of *DHX38* (sgRNA_10966) and *MAT2A* (sgRNA_02249) using the tools transIndel and Genesis-Indel, which are specifically designed to identify indels in the unmapped read fraction of samples^60,61^.

### Analysis of scRNA-seq data from human coronary arteries

Single-cell gene expression matrix from human right atherosclerotic coronary arteries (three male and one female donors), was downloaded from NCBI GEO (GSE131780, https://www.ncbi.nlm.nih.gov/geo/query/acc.cgi?acc=GSE131780). The data was re-analyzed using the Seurat package in R with a standard single-cell clustering pipeline. Gene expression data was normalized using the SCTransform function from Seurat (v.3.2.3), regressing out the percentage of mitochondrial gene expression. Principal components analysis was performed, followed by dimensional reduction with Uniform Manifold Approximation and Projection (UMAP) using the first 20 principal components as input. Gene expression was visualized on the first two UMAP dimensions using the kernel density function (plot_density) from the Nebulosa package (v.0.99.92)^62^ for endothelial cell marker and candidate genes.

### Statistics and data analysis

Unless noted otherwise, we performed all data and statistical analyses in R (v.3.6.0) using Rstudio. We ran our analyses on a high performance computing cluster (Beluga) from Calcul Quebec/Compute Canada. For MAGECK variant-level analyses, permutation-based FDR of ≤10% were considered significant. For RNA-seq analysis, genes with a Benjamini-Hochberg adjusted P-value in DESeq2 ≤0.05 were considered significant^50^.

## Acknowledgements

This research was enabled in part by support provided by Calcul Quebec (https://www.calculquebec.ca/en/) and Compute Canada (www.computecanada.ca). We thank John D. Rioux and Catherine Martel for flow cytometry and FACS analysis. We thank Éric Thorin, Mike Sapieha, Dan Bauer and Luca Pinello for providing comments on an earlier version of this manuscript. Finally, we thank Génome Québec and the Center for Applied Genomics (TCAG) in Toronto for performing next-generation DNA sequencing for this project.

## Data availability

The data discussed in this publication have been deposited in NCBI’s Gene Expression Omnibus^63^ and are accessible through GEO Series accession number GSE165925 (https://www.ncbi.nlm.nih.gov/geo/query/acc.cgi?acc=GSE165925).

