## Supplementary Figures for "CRISPR perturbations at many coronary artery disease loci impair vascular endothelial cell functions"

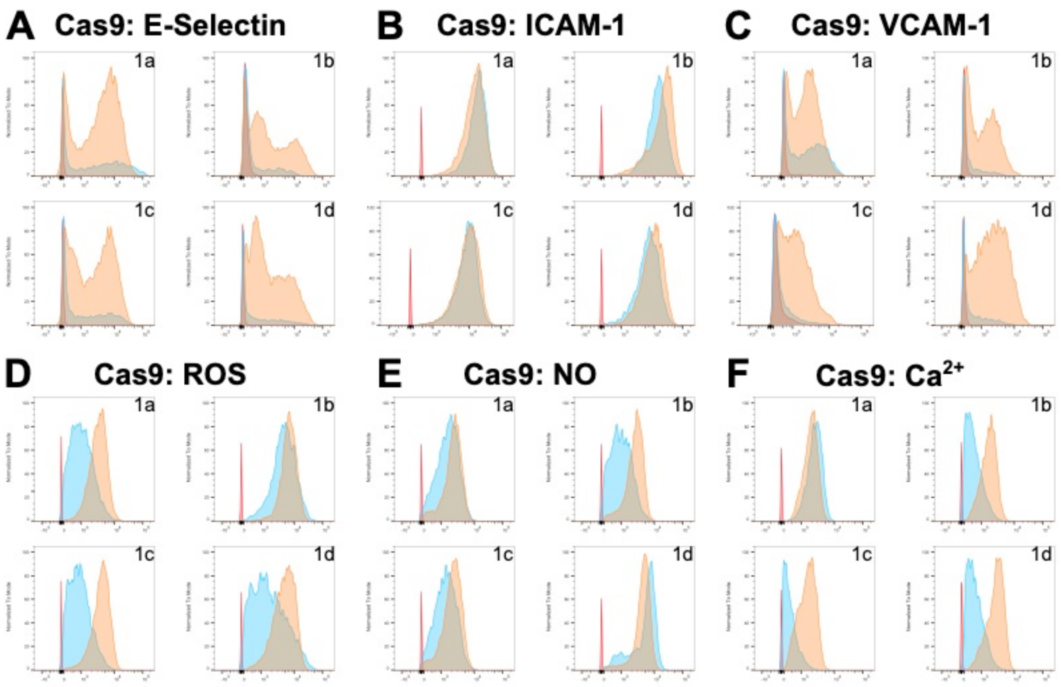


**Supplementary Figure 1.** Flow cytometry profiles for all six endothelial cell readouts with Cas9.

For each experiment, we show a representative figure of the flow cytometry profiles obtained for each of the four lentiviral batches (1a-1d) used in our experiments. Red: Non-infected, non-stained cells; Blue: Non-infected, stained cells; Orange: Infected and stained cells.

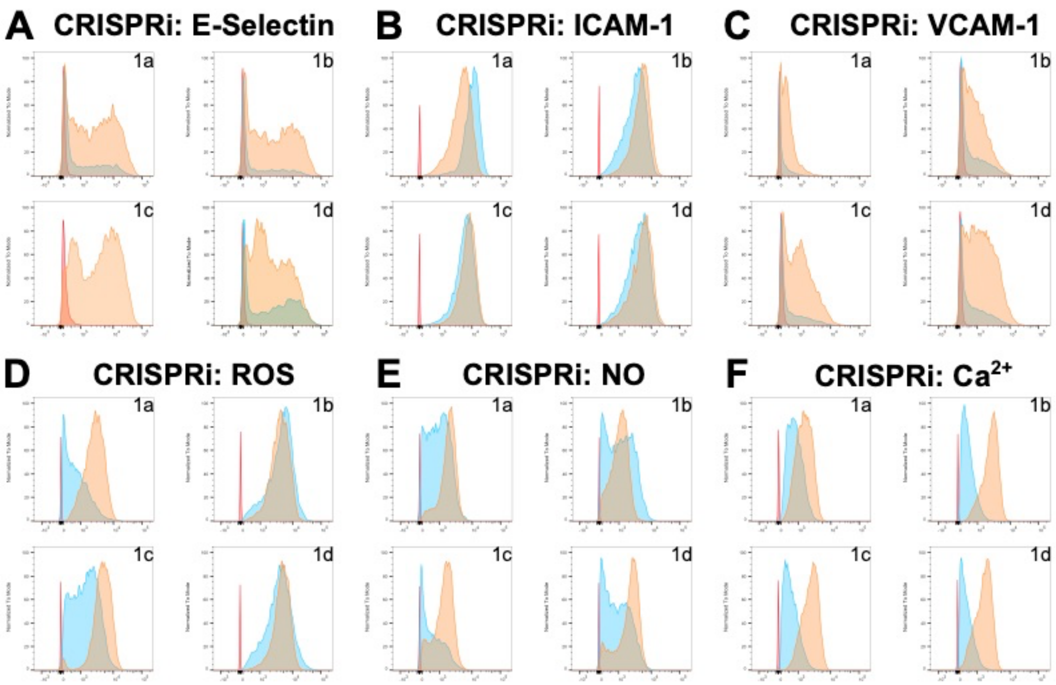


**Supplementary Figure 2.** Flow cytometry profiles for all six endothelial cell readouts with CRISPRi.

For each experiment, we show a representative figure of the flow cytometry profiles obtained for each of the four lentiviral batches (1a-1d) used in our experiments. Red: Non-infected, non-stained cells; Blue: Non-infected, stained cells; Orange: Infected and stained cells.

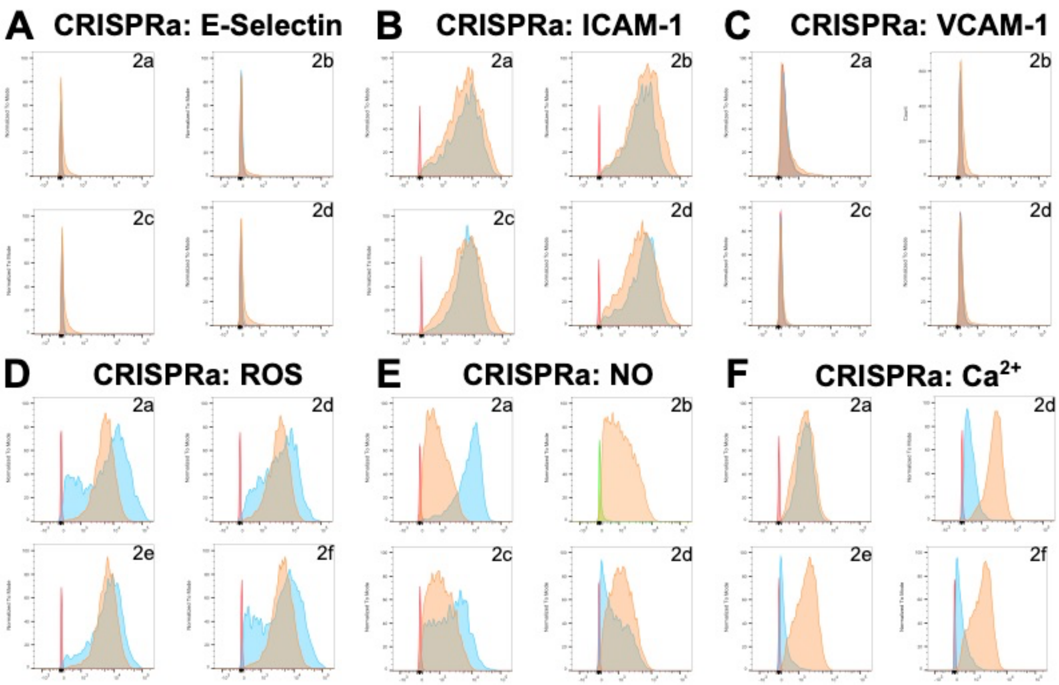


**Supplementary Figure 3.** Flow cytometry profiles for all six endothelial cell readouts with CRISPRa.

For each experiment, we show a representative figure of the flow cytometry profiles obtained for each of the four lentiviral batches (1a-1d) used in our experiments. Red: Non-infected, non-stained cells; Blue: Non-infected, stained cells; Orange: Infected and stained cells.

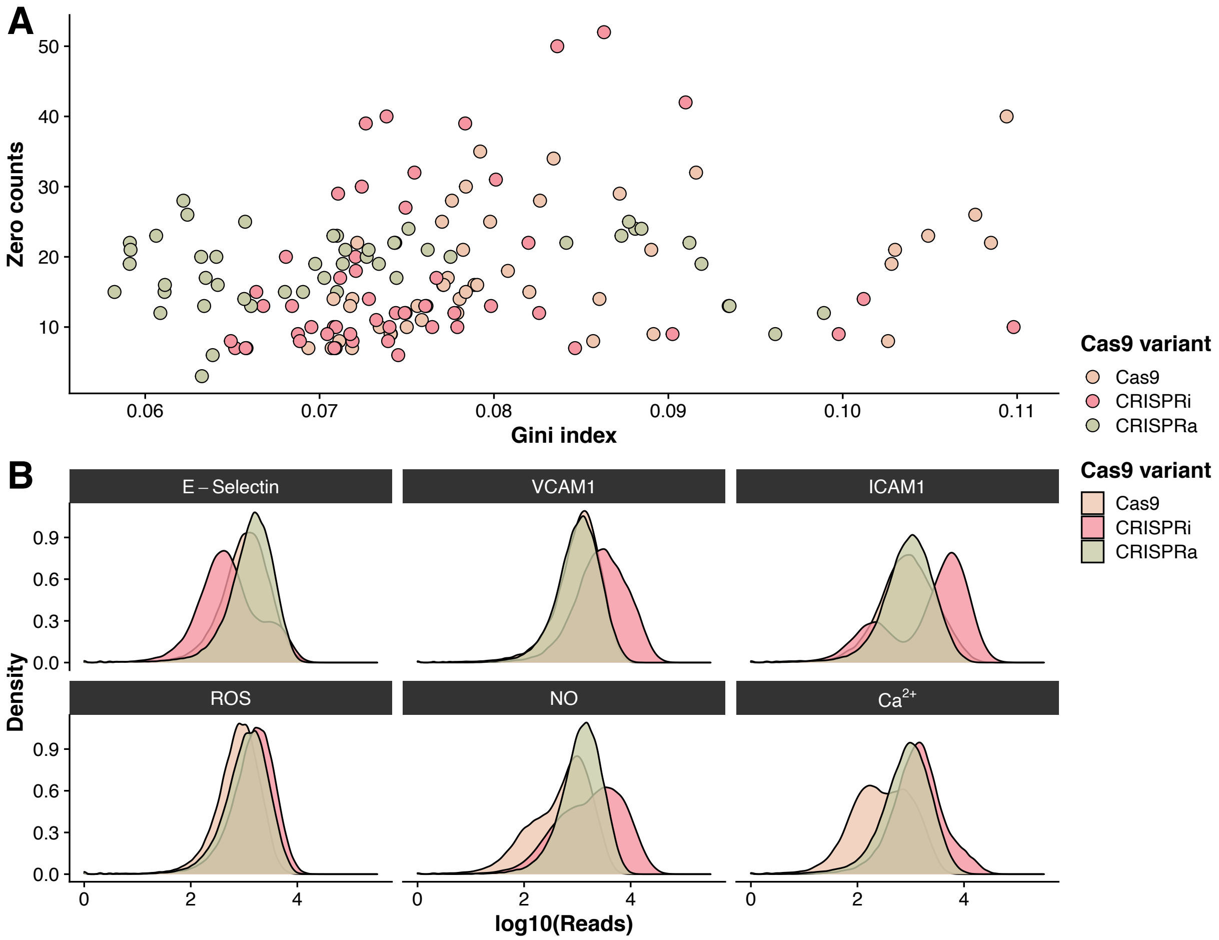


**Supplementary Figure 4.** Quality-control metrics of sequenced lentiviral libraries.

(**A**) The Gini index is a measure of sgRNA diversity across a sample. A low Gini index indicates that all sgRNAs are equally represented in the sequenced data. The data is stratified (color-coded) by Cas9 proteins. For the sequence data across all experiments, the Gini index is below the recommended threshold (Gini index <0.2) [1]. The *y*-axis denotes the number of sgRNAs with 0 reads for each experiment.

(**B**) Density distribution of reads per sgRNA for all endothelial phenotypes and Cas9 proteins (the x-axis is read depth on a $log_{10}$ scale).

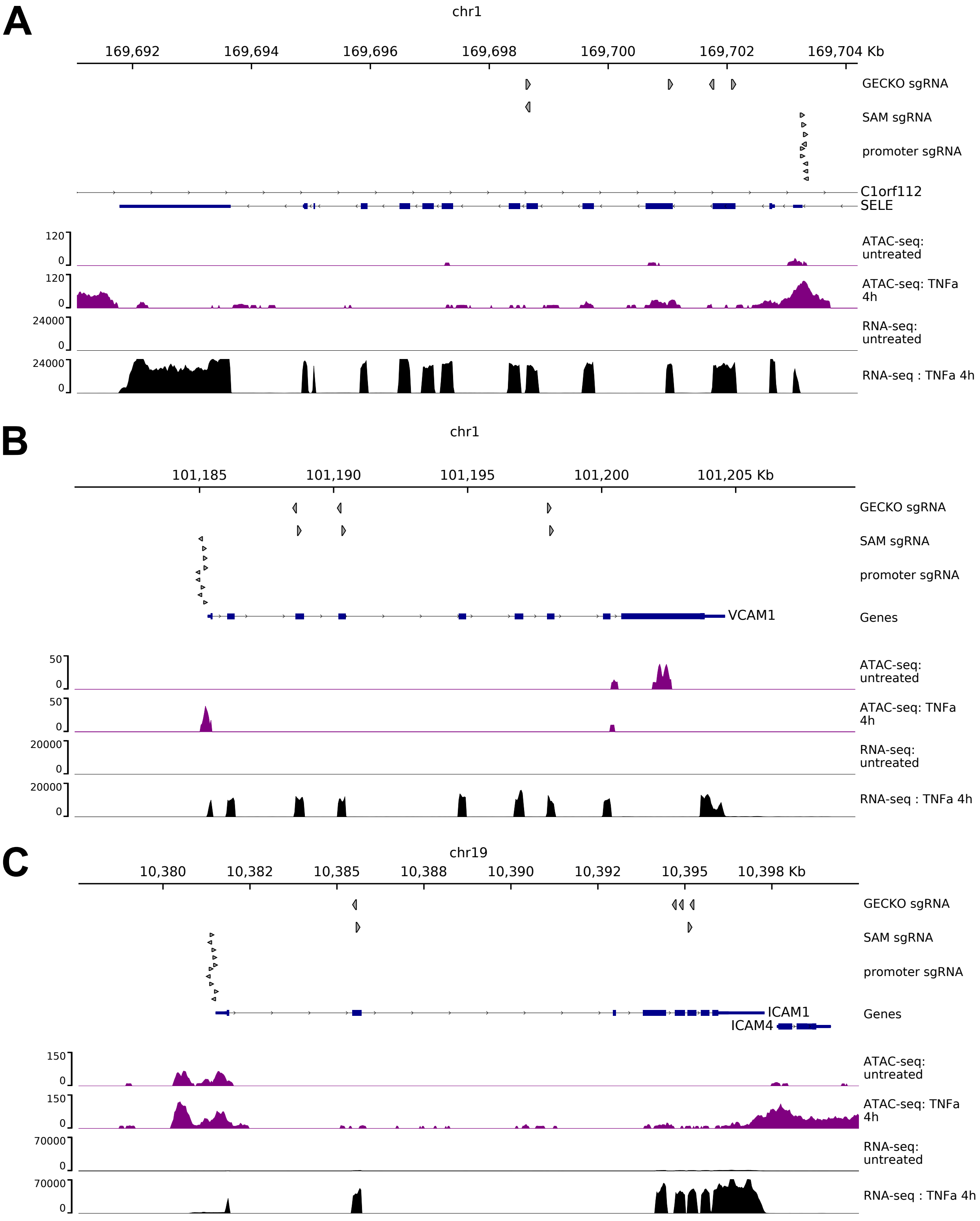


**Supplementary Figure 5.** Graphical representation of three genes that encode adhesion molecules important for monocyte rolling and attachment at the onset of atherosclerosis.

Locus views for adhesion molecule positive control genes (**A**) *SELE* (E-selectin), (**B**) *VCAM1* and (**C**) *ICAM1*. We also represent the position of the sgRNAs that were tested in the pooled CRISPR screens (GECKO, coding sequences; SAM and promoter, regulatory sequences), as well as RNA-seq and ATAC-seq data in teloHAEC that are un-stimulated or activated for 4 hours with TNFα [2].

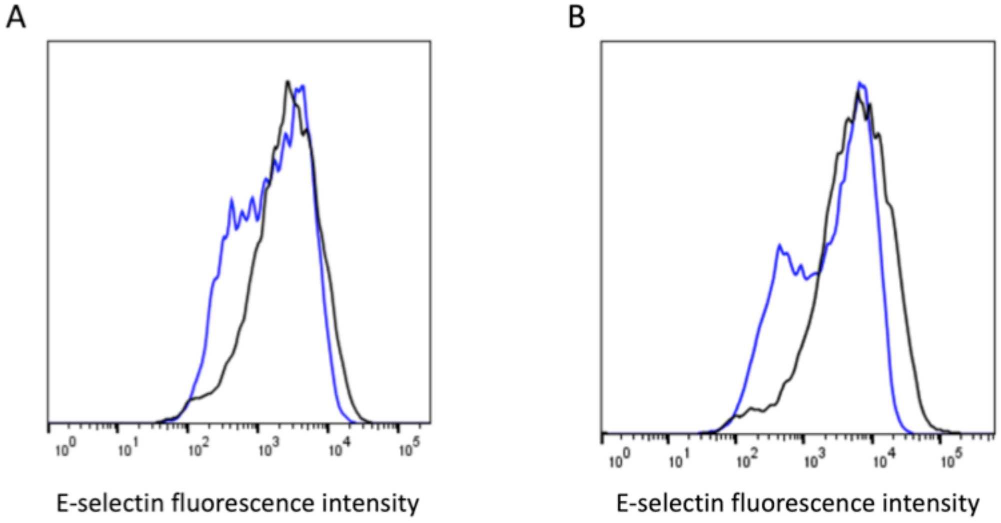


**Supplementary Figure 6.** Flow cytometry experiments for E-selectin using nucleofection of ribonucleoprotein (RNP) complexes containing Cas9 and sg_10966 (rs2074626 in *DHX38*).

Results are shown for two independent biological replicates . At the 10% bottom fraction threshold for E-selectin levels in the control experiments (empty nucleofection, black), we find 24% (panel A) and 31% (panel B) of cells following Cas9 RNP targeting rs2074626 in *DHX38*. This result confirms that *DHX38* inactivation reduces E-selectin presentation at the cell membrane.

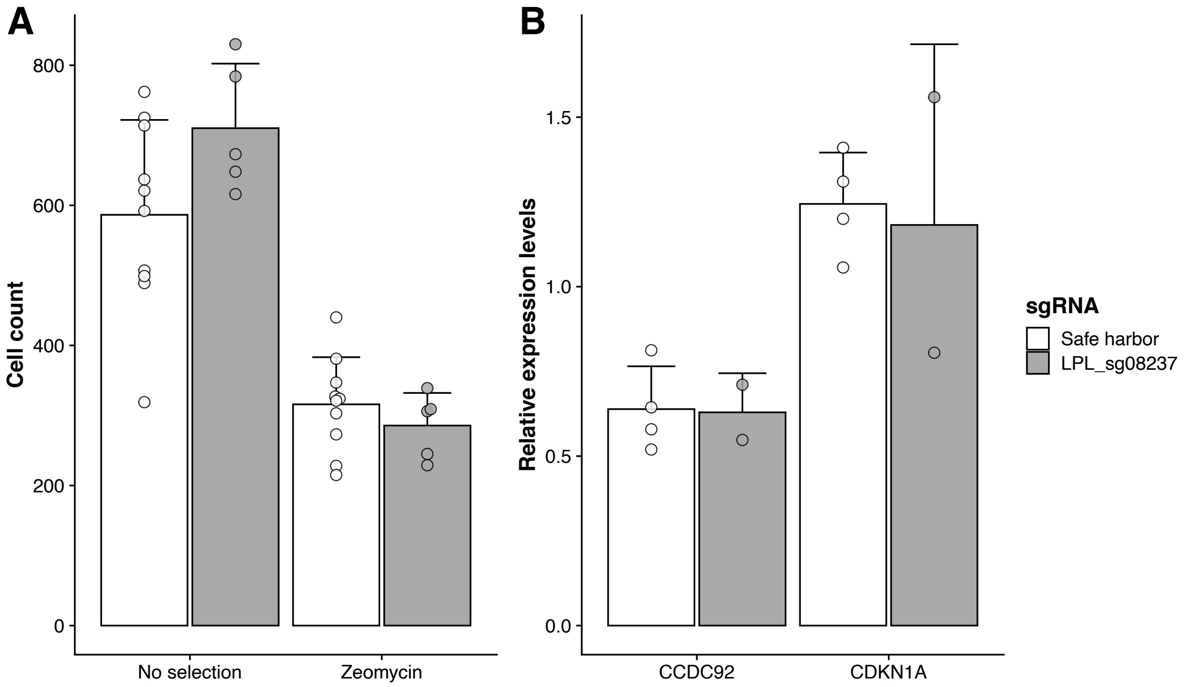


**Supplementary Figure 7.** CRISPRa experiment with a silent sgRNA at the *LPL* locus does not influence cell proliferation nor the expression of senescence marker genes.

(**A**) teloHAEC that express dCas9-VP64 were infected with a lentivirus that carries a sgRNA that targets rs1441755 at the LPL locus. This sgRNA was silent in all our pooled CRISPR screens for all six endothelial phenotypes tested. In the absence or presence of antibiotic selection (Zeocin), LPL_sg08237 does not affect cell proliferation. Cell counts are mean ± standard deviation of 10 and 5 images for the safe harbor and LPL_sg08237 sgRNAs, respectively. The differences are not significant (Student's *t*-test $P$>0.06).

(**B**) Expression of *CCDC92* and *CDKN1A* in teloHAEC that express dCas9-VP64 (with zeocin selection). Results are mean ± standard deviation from 4 and 2 experiments for the safe harbor and LPL_sg08237 sgRNAs, respectively. The differences are not significant (Student's *t*-test $P$>0.6).
